# A UNIQUOME BASED METHOD FOR THE PROTEIN IDENTIFICATION BY MASS SPECTROMETRY

**DOI:** 10.1101/2025.10.08.681213

**Authors:** Pierros Vasileios, Kontopodis Evangelos, Stravopodis J. Dimitrios, Tsangaris Th. George

## Abstract

Protein identification by mass spectrometry is a pivotal step in proteomics. Numerous methods have been developed to securely and effectively identify proteins derived from experimentally detected peptides by mass spectrometry. The dominant approach is based on the assumption that each experimentally identified peptide can be matched with a peptide included in a database of peptide sequences, generated by in silico digestion of proteins with a specific proteolytic enzyme. In this way, the protein containing the peptide can be identified. In a more advanced approach, the proteins and their in silico–digested peptides in the database are transformed into theoretical mass spectrometry spectra, and search engines match the experimentally obtained spectra to these theoretical spectra generated from protein and peptide sequences. We developed an alternative method for protein identification using Core Unique Peptides (CrUPs) and the Uniquome, termed as **Uniquome-Based Protein Identification Method (UB-PIM)**. According to this method, instead of searching for peptides in the database of *in silico*–digested peptides, we search for CrUPs within the experimentally obtained peptides by mass spectrometry. If a peptide contains at least one CrUP, it can be directly correlated to the protein from which the CrUP is derived. Because of the unique nature of CrUPs, peptides obtained by MS can securely and uniquely identify the protein of origin. This provides a reference space in which even single-peptide identifications can achieve high specificity, reducing the ambiguity caused by shared or homologous sequences and improving the interpretability of MS data. Furthermore, UB-PIM can be applied to any type of peptide and is effective with both Data-Independent Acquisition (DIA) and Data-Dependent Acquisition (DDA) approaches, as well as with top-down and bottom-up proteomics. This allows confident protein identification from minimal evidence, expands the scope of detectable proteins, and remains computationally efficient, rapid, and universally applicable.

## 1. Introduction

Proteomics is a set of complex methods and technologies aimed at identifying, cataloging, and studying the total protein content of a biological material. The conventional approach includes the separation of proteins in the biological sample, their digestion by proteolytic enzymes, the analysis of the resulting peptides by mass spectrometry, protein identification using bioinformatics tools, systematic entry of the results into databases, and finally their processing and evaluation. The introduction of mass spectrometry (MS) proved pivotal, and the development of MS-based proteomics has profoundly transformed the ability to identify and quantify proteins in complex biological samples (1).

At its core, proteomic workflows employ either “bottom-up” or “top-down” approaches—digesting proteins into peptides or analyzing intact proteins directly, respectively. MS methods can be broadly divided into Data-Independent Acquisition (DIA) and Data-Dependent Acquisition (DDA) techniques (2, 3). DIA is the MS technique primarily used in proteomics to identify and quantify proteins in complex biological samples. It systematically fragments all ions within predefined mass-to-charge (m/z) windows across the entire m/z range, unlike DDA, where only a subset of ions is selected for fragmentation. In both techniques, protein identification relies heavily on matching observed mass-to-charge (m/z) spectra to peptide/protein sequences through computational search engines and scoring systems, aided by statistical validation techniques such as false discovery rate (FDR) control (2–4).

Briefly, protein identification in a proteomics workflow is based on a FASTA-format file containing peptide sequences, generated by in silico digestion of proteins with a specific proteolytic enzyme (commonly trypsin, but also chymotrypsin, etc.). To generate the FASTA file, all reviewed and unreviewed proteins of a given species (e.g., *Homo sapiens*) from the UniProt database are digested in silico according to the rules of the selected enzyme, and the resulting theoretical peptides are compiled into the FASTA file (5). For protein identification, each experimentally observed peptide by MS is matched with a peptide contained in the FASTA file, thereby identifying the protein in which the peptide occurs (6).

In a more advanced approach, proteins and their *in silico*–digested peptides included in the FASTA file are transformed into theoretical MS spectra, and search engines (such as Mascot, MaxQuant, MSFragger, or Comet) match the experimentally obtained spectra to these theoretical spectra generated from protein and peptide sequences (7, 8).

Despite their widespread use, these methods have important limitations. For reliable protein identification, the analysis of at least two peptides per protein is typically required, and many peptides identified by the mass spectrometer do not ultimately lead to unambiguous protein characterization and are therefore rejected during bioinformatics processing. Increasing the number of search engines and integrating diverse algorithmic approaches can marginally improve protein identification coverage, but such gains are often limited by computational burden and diminishing returns (9). These weaknesses of current methods have highlighted the need for a new approach to protein identification. Such an approach is based on the assumption that the amino acid sequence of each protein should contain at least one peptide whose sequence is absolutely unique to the proteome of the organism to which it belongs, thereby enabling differential and unambiguous characterization of the protein (10– 12).

In this context, the concept of the **Uniquome** offers a groundbreaking strategy, originally introduced for constructing a proteomic atlas for each organism by cataloging peptides whose amino acid sequences are absolutely unique to that organism’s proteome. Within the Uniquome, two novel peptide entities were defined: a) the Core Unique Peptide (CrUP), which is a peptide sequence present specifically and exclusively in only one protein of a given proteome, and which also represents the minimal amino acid length necessary for unique identification; and b) the Composite Unique Peptide (CmUP), an extended sequence entity formed by the linear, sequential, or overlapping unification of two or more CrUPs (10, 11). Thus, a typical CrUP is the shortest amino acid sequence found exclusively in a single protein within a proteome, ensuring that the sequence does not occur in any other protein of the examined proteome. The term “core” emphasizes that this peptide is minimal in length yet sufficient to uniquely represent its source protein (13).

Analysis of the human proteome revealed that 99.2% of reviewed proteins contain CrUPs, suggesting that leveraging these peptide entities could dramatically enhance identification specificity—even in cases supported by only a single peptide. This challenges the conventional requirement for multi-peptide evidence. By computing and indexing these unique sequences, the Uniquome provides a reference space in which single-peptide identifications can achieve high specificity, thereby reducing ambiguity caused by shared or homologous sequences and improving the interpretability of MS data (12, 13).

Based on this framework, we developed an MS-based protein identification method in which the counterpart of the FASTA file used in conventional approaches is replaced with a file containing the CrUPs of the Uniquome of the studied organism. In the present study, this method was applied to the identification of *Homo sapiens* proteins included in the human proteome. The obtained results indicate that the Uniquome could transform MS-based protein identification in both DIA and DDA approaches by removing ambiguity at the sequence level, enabling confident identification from minimal evidence, and expanding the scope of detectable proteins—all while being computationally efficient, rapid, and universally applicable.

## 2. Materials

### 2.1. Databases used

The following databases were used:

- UniProt (www.uniprot.org) (Release 2025_03);
- Unipept (https://unipept.ugent);
- InterPro (https://www.ebi.ac.uk/interpro) (Release 97.0, 09 Nov.2023);
- National Center for Biotechnology Information (https://www.ncbi.nlm.nih.gov);
- Kyoto Encyclopedia of Genes and Genomes (https://www.genome.jp/kegg/);
- Japan ProteOme STandard Repository (https://repository.jpostdb.org);
- Trans-Proteomic Pipeline (TPP) (http://www.tppms.org)

## 3. Methods

### 3.1. Uniquome and Uniquome (+) databases

In the latest release of the human proteome in the UniProt database (Release 2025_3), 20,420 reviewed proteins and 184,785 unreviewed proteins were included [14]. The total set of 20,420 reviewed human proteins was analyzed for the presence of the novel CrUP and CmUP peptide entities. Following the procedure described previously [13], a total of 7,279,076 peptides ranging from 4 to 100 amino acids (aa) in length were characterized as CrUPs in 20,251 (99.16%) human proteins (Table 1), whereas 170 proteins (0.82%) did not contain any UP species of the same length (4–100 aa). Further analysis revealed that the 7,279,076 CrUPs could generate 67,921 CmUPs. The overall density of the human proteome in CrUPs is 64%, indicating that 64 CrUPs are generated per 100 aa, while the coverage of the human proteome by UP species is 93% (Table 1).

**TABLE 1.** Uniquome and Uniquome (+) analysis.

|  | **Uniquome** | **Uniquome (+)** |
| --- | --- | --- |
| **Proteins included** | 20,420<br>(Only reviewed) | 20,859<br>(Reviewed + Unreviewed) |
| Core Unique Peptides | 7,279,076 | 7,279,514 |
| Composite Unique Peptides | 67,921 | 67,932 |
| Density of CrUP | 64% | 64% |
| Density of CmUP | 0.6% | 0.6 |
| Coverage | 93% |  |
| Proteins with Unique Peptides | 20,251 | 20,385 |
| Proteins without Unique Peptides | 170 | 474 |

For protein identification by CrUPs, the construction of the Human Uniquome database is crucial. In our previous studies, the Uniquome database was constructed exclusively from reviewed proteins in UniProt. To expand this database, we examined the possibility of including unreviewed proteins from UniProt. Analysis of these proteins revealed several cases: (a) proteins with identical sequences to reviewed proteins but with different accession numbers. For example, the reviewed protein SAMP_Human (P02743, Serum Amyloid P-component) has an identical amino acid sequence to the unreviewed protein V9HWP0_Human (V9HWP0, Pentraxin Family member) (Sup. Fig. 1A). In this case, the CrUPs of SAMP_Human lose their uniqueness, as they also belong to V9HWP0_Human. (b) Fragments of reviewed proteins, which share identical sequence segments with the complete protein but are assigned different accession numbers. For instance, the reviewed protein VW5B1_Human (Q5TIE3, von Willebrand factor A domain-containing protein 5B1) has a length of 1220 amino acids, whereas the unreviewed protein E9PP07_Human (E9PP07, von Willebrand factor A domain-containing 5B1) is an 82-aa fragment corresponding to the N-terminal region of VW5B1_Human (Sup. Fig. 1B). In this case, the CrUPs located within the first 82 amino acids of VW5B1_Human lose their uniqueness, as they also belong to E9PP07_Human.

To expand the Human Uniquome database while accounting for such cases, we constructed the Uniquome (+) database. This extended version was designed to include only unreviewed proteins that do not “overlap” with any reviewed proteins. Specifically, unreviewed proteins that did not contain any CrUPs of the reviewed proteins in their sequence were isolated. These proteins, lacking CrUPs derived from reviewed proteins, indicate that they share no amino acid sequence with reviewed proteins and are not fragments of them. We identified 439 such unreviewed proteins. These proteins were added to the 20,420 reviewed proteins, creating a new dataset of 20,859 proteins (Table 1). In this expanded set, CrUPs and CmUPs were re-calculated, resulting in 7,279,514 CrUPs and 67,932 CmUPs, respectively, which together constitute the Uniquome (+) (Table 1).

### 3.2.The. Uniquome Based-Protein Identification Method (UB-PIM)

We developed a protein identification algorithm based on the Uniquome and Uniquome (+) databases, which checks whether an MS-identified peptide contains at least one CrUP. If the check is positive, the peptide is directly associated with the corresponding protein from which the CrUP is derived. Since each CrUP is unique to its source protein, any MS-identified peptide containing it can unambiguously and reliably identify that protein. In other words, the presence of a CrUP within a peptide ensures that the peptide uniquely identifies a single protein.

The developed method, named as **Uniquome-Based Protein Identification Method (UB-PIM)**, has been implemented on the Uniquome.com website. Figure 1, shows a screenshot of the Uniquome.com homepage, where protein identification via the developed method is introduced as *“Protein identification (UB-PIM)*.*”* Selecting this option leads the user to the main interface of the UB-PIM method (Figure 2). On this screen, required and optional fields are clearly indicated by an asterisk (*).

**Figure 1:**
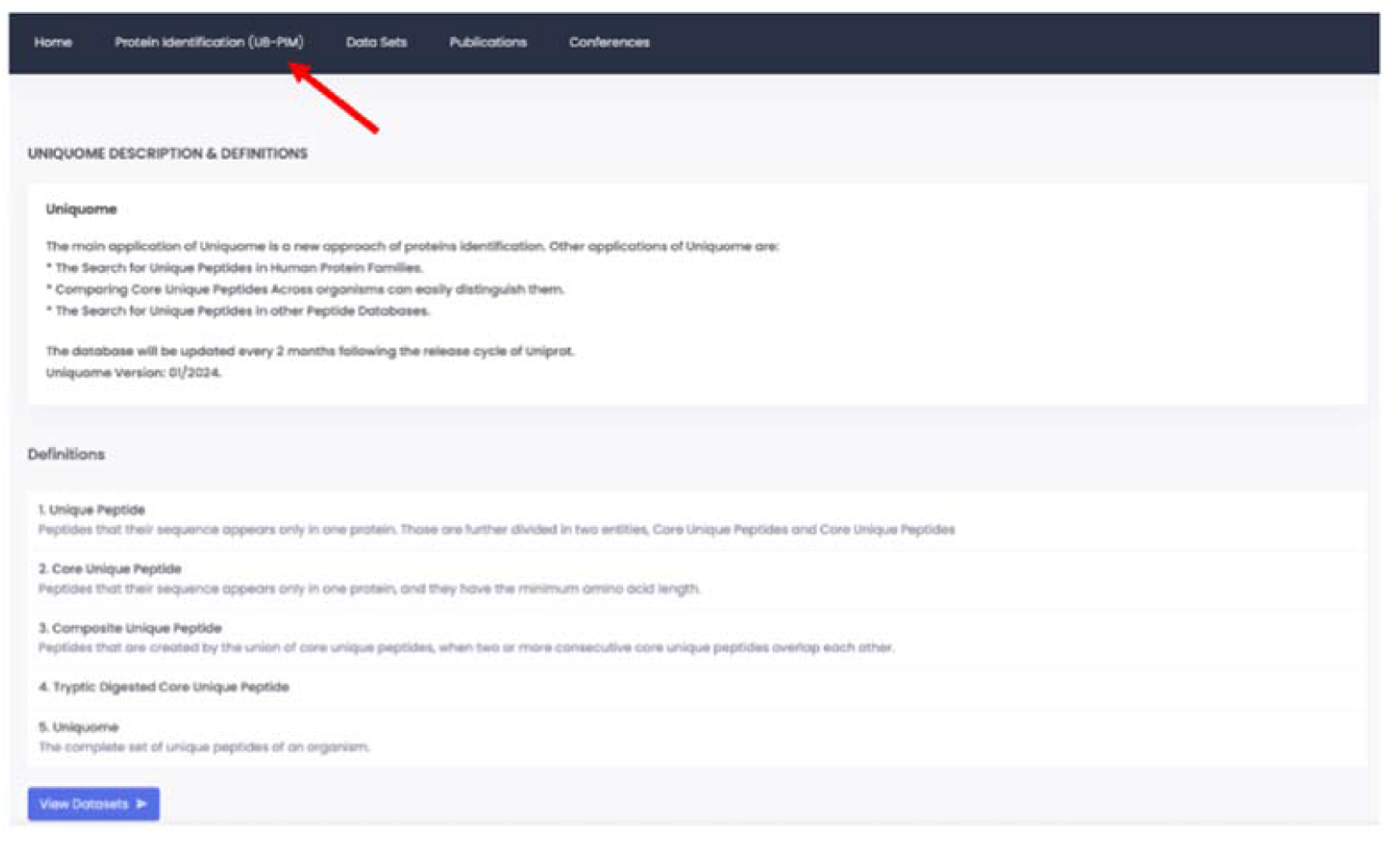
The Uniquome.com homepage, where protein identification via the developed method is introduced as *“Protein identification (UB-PIM)”*. The position in the page is marked by the red arrow.

**Figure 2:**
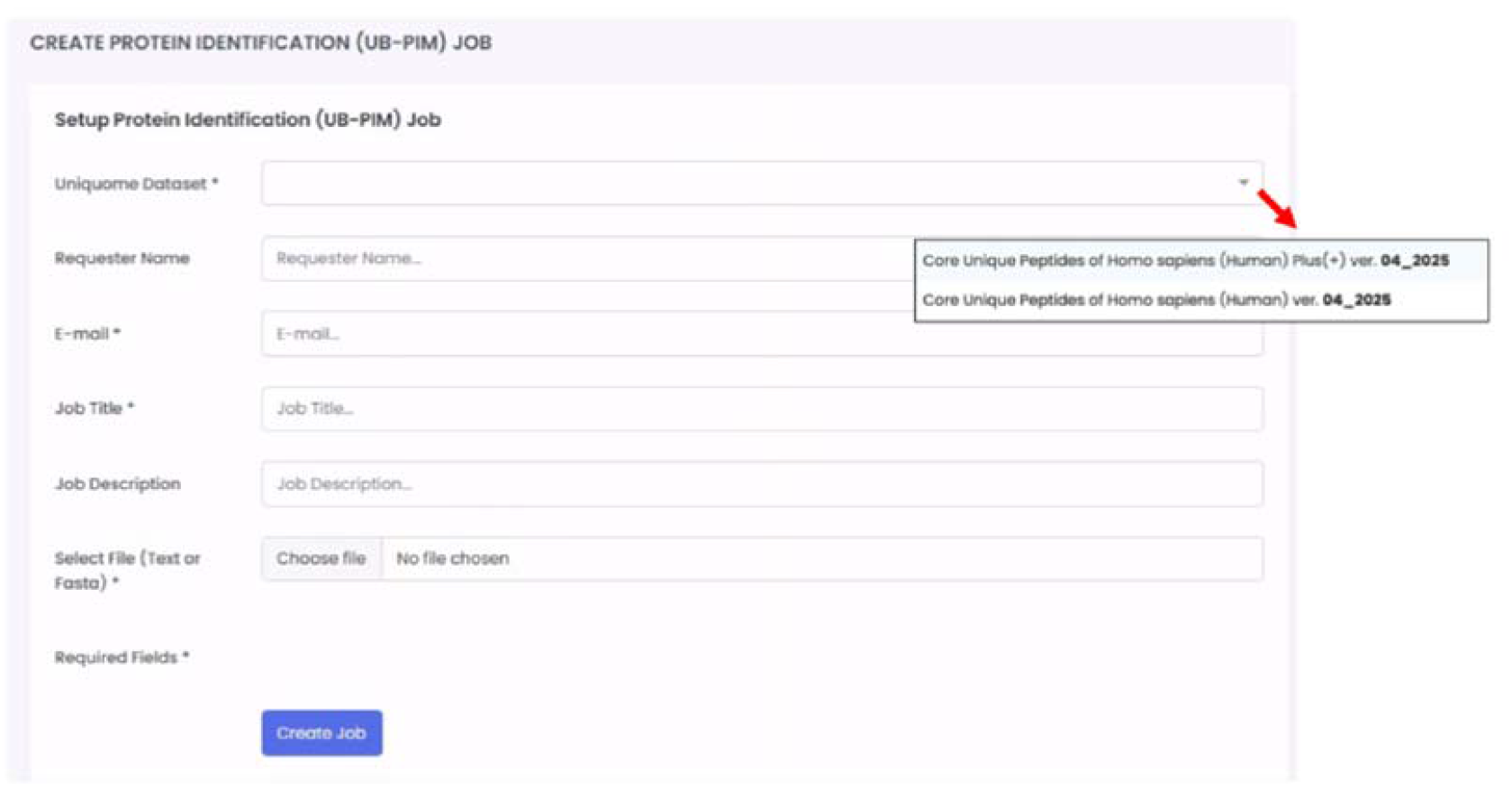
The main interface of the UB-PIM method after the Uniquome.com homepage. Required and optional fields are indicated by an asterisk (*). Red arrow indicate that in the required position “Uniquome Dataset” the user could select the Core Unique Peptides of Homo Sapiens (Human) Plus (+) or the Core Unique Peptides of Homo Sapiens (Human).

In the first required field, the user must select which database will be used for protein identification. Two options are available: the latest version of the Human Uniquome [appearing as *Core Unique Peptides of Homo sapiens (Human) ver. 4_2025*] or the latest version of the Uniquome (+) [appearing as *Core Unique Peptides of Homo sapiens (Human) plus (+) ver. 4_2025*]. The second required field is the user’s email address, to which the results will be sent. The third required field is the “Job Title,” where the user provides a descriptive name for the task. Finally, under “Select File,” the user uploads the file containing the MS-derived peptides to be analyzed. This file may be either a plain text file (with one peptide per line) or a FASTA-format file. In the case of a text file, peptides are automatically numbered (e.g., 000001), whereas in a FASTA file, peptides can be grouped under user-defined names. For demonstration purposes, a list of 30 peptides in text format was used to perform the method (Table 2). Optional fields allow the user to enter their name and a more detailed description of the submitted task, but these are not mandatory.

**Table 2.**
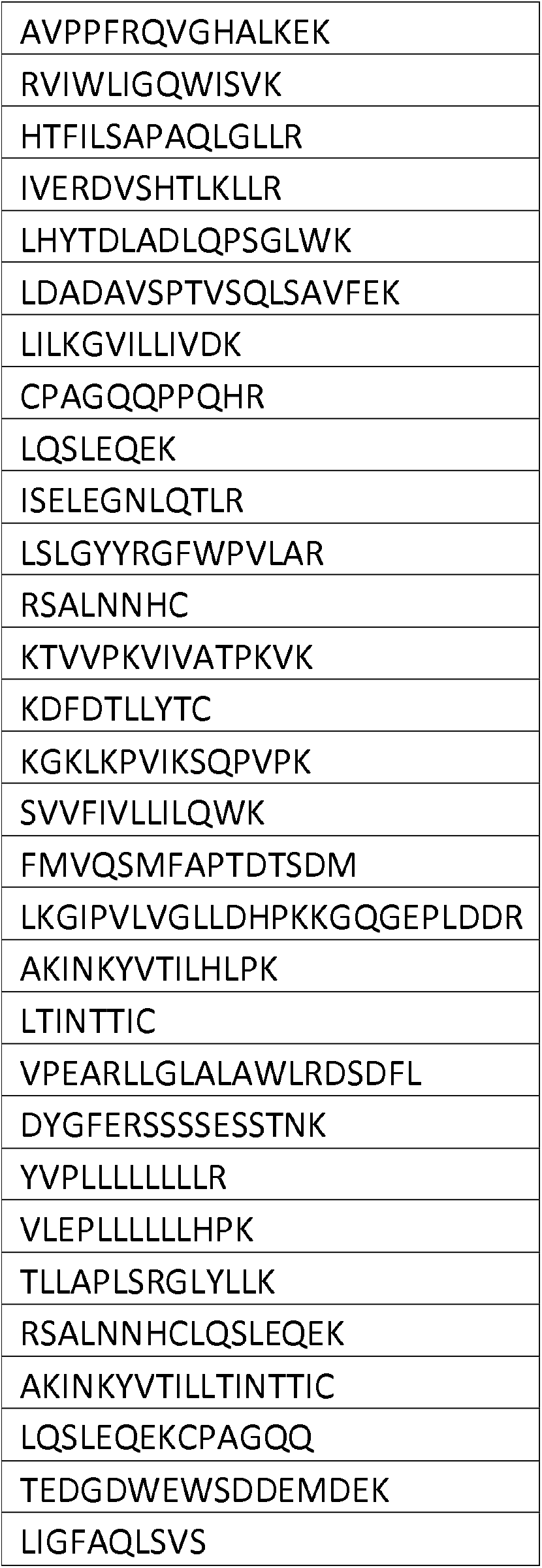
Peptides.

After all required fields are completed, clicking “Create Job” initiates the method. UB-PIM is executed asynchronously at scheduled intervals (every few hours). The process begins by retrieving all proteins and their corresponding Core Unique Peptides (CrUPs) from the selected database. Each user-provided peptide is then checked to determine whether it contains one or more CrUPs from a given protein. A peptide may match multiple CrUPs from the same protein. However, if a peptide does not contain any CrUP or contains CrUPs from more than one protein, it is flagged as *“Unable to identify protein”* and excluded from the results matrix. After processing all peptides, the method compiles the distinct set of proteins identified—while accounting for the fact that a single protein may be matched by multiple peptides.

### 3.3. UB-PIM results interpretation

The results are exported as an Excel file, which is automatically sent to the email address specified by the user. The resulting Excel file contains five worksheets (Figure 3):

1. **Job Details** – Provides information about the submitted job.
2. **Results** – Lists all peptides included in the user file, along with the proteins they identify, as well as peptides unable to identify a protein.
3. **Identified Proteins** – Summarizes the proteins identified from the peptides in the input file.
4. **CrUPs Matches** – Provides detailed information on the Core Unique Peptides (CrUPs) matched to each user peptide.
5. **Findings Matrix** – Displays all peptides that identify a protein, with an “X” marking the intersection between peptides and their corresponding proteins.

**Figure 3:**
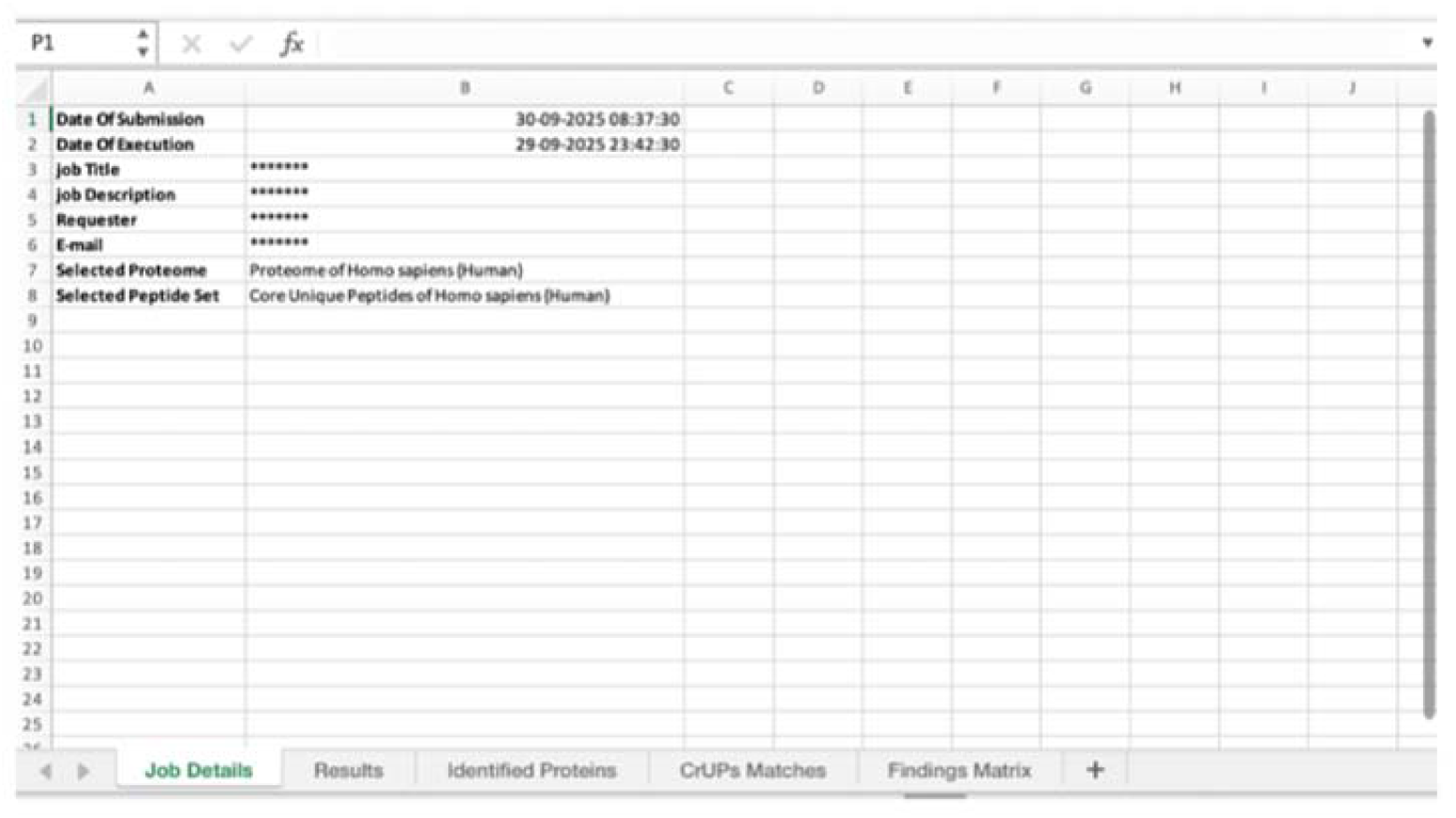
The Excel file which the user receives automatically after applying the file containing the MS-derived the peptides to be analyzed by UB-PIM. The Excel file includes five worksheets as shown in the bottom of the excel file.

In our example, the *Results* worksheet includes all peptides listed in Table 2, along with the corresponding protein identified by each peptide after applying the developed method. Four peptides were classified as *“Unable to identify protein”* and are highlighted in red (Figure 4).

**Figure 4:**
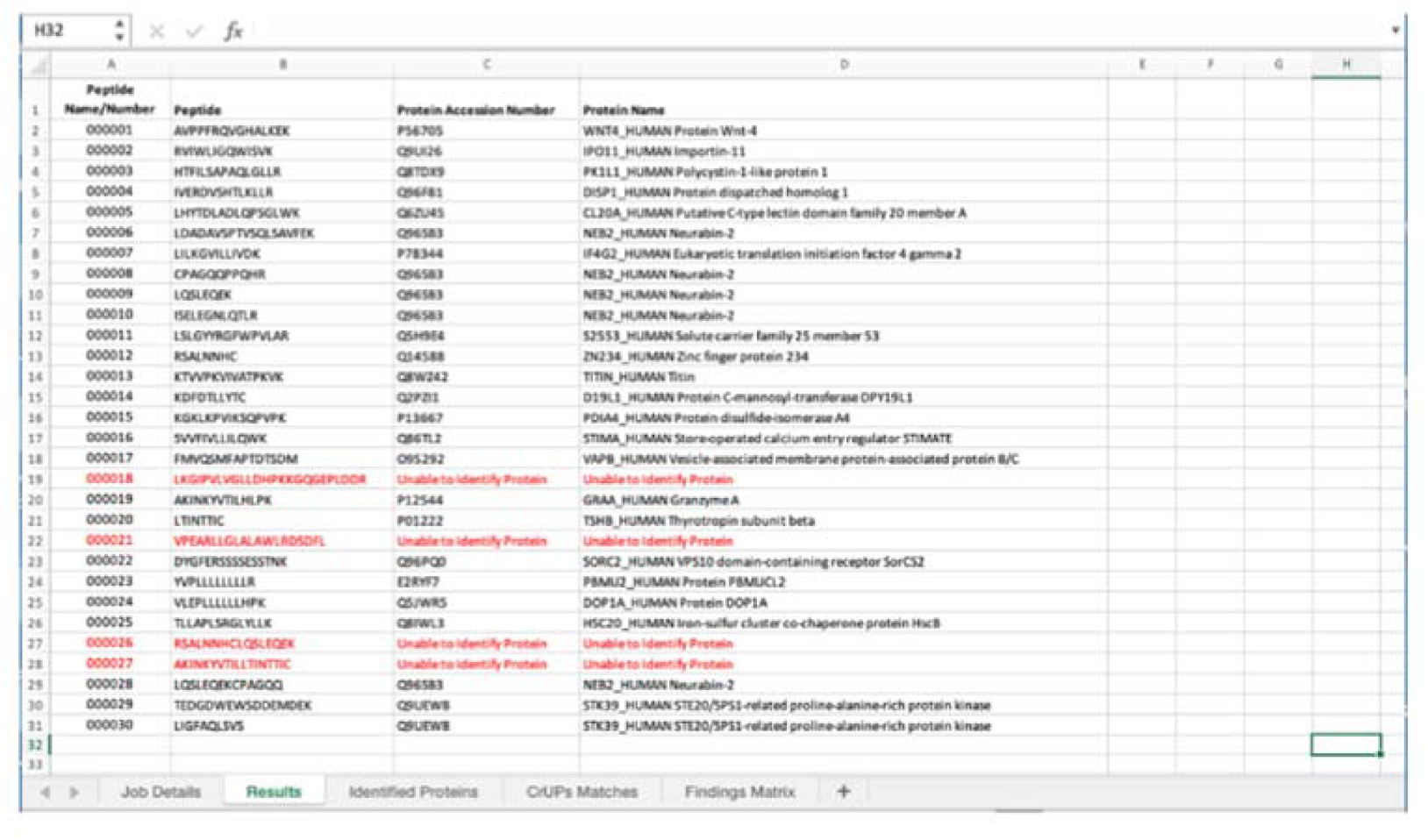
The “Results” worksheet. Lists all peptides included in the user file, along with the proteins they identify. Peptides unable to identify a protein appeared in red color marked by the “Unable to Identify Protein”.

The next worksheet, *Identified Proteins*, provides a summary of proteins identified by the uploaded peptides and their associated CrUPs (Figure 5). In this example, 21 proteins were identified by 26 of the 30 uploaded peptides, while 4 peptides did not identify any protein. The first column lists the UniProt accession number of each identified protein, and the second column provides its full protein name. The third column shows the number of peptides from the input set that contributed to the identification of each protein. For example, 16 proteins were identified by a single peptide, one protein by 2 peptides, and one protein by 5 peptides. The fourth column lists the number of identified CrUPs per protein, while the fifth column shows the total number of CrUPs present in each identified protein. The final column contains a comma-separated list of all matched CrUPs for each protein.

**Figure 5:**
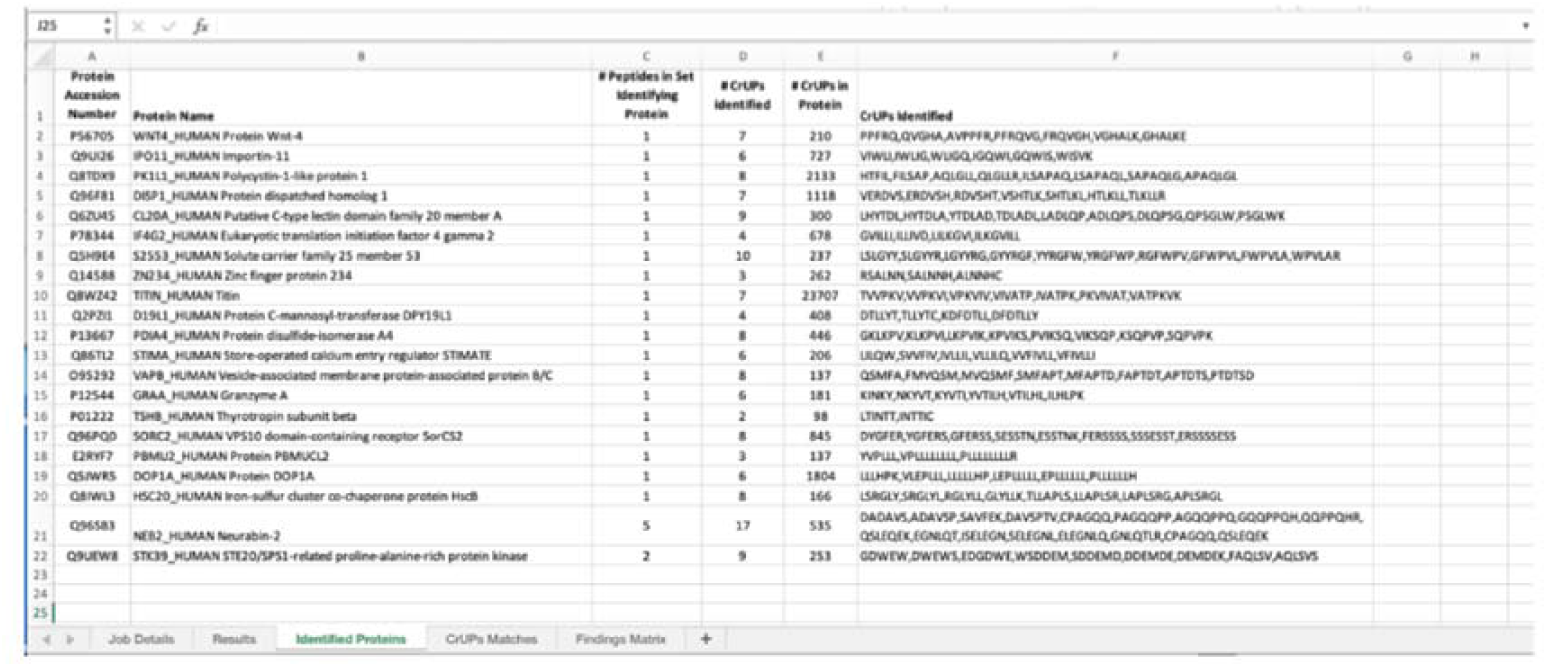
The “Identified Proteins” worksheet. The identified proteins by the uploaded set of peptides, the number of CrUPs and a comma-separated list of all matched CrUPs for each protein are shown.

The *CrUPs Matches* worksheet presents the detailed CrUP information underlying protein identification (Figure 6). Specifically, it shows the CrUP(s) included in each peptide, the protein of origin, the position of the CrUP within the peptide, and the position of the CrUP within the corresponding protein sequence.

**Figure 6:**
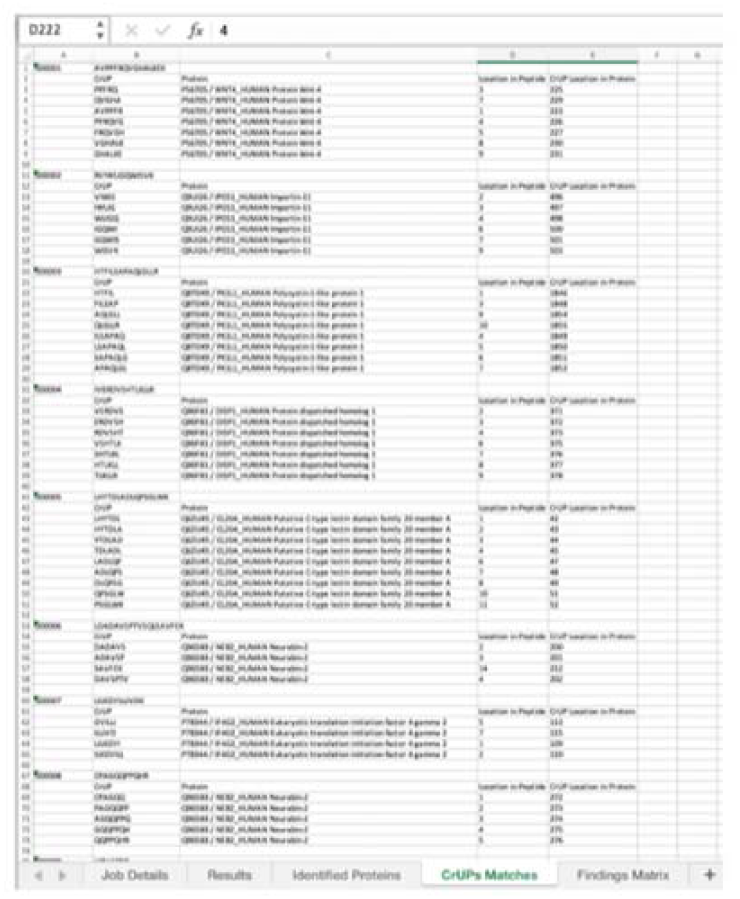
The “CrUPs Matches” worksheet. Detailed information on the CrUPs matched to each uploaded peptide together with the identified protein and the location of each CrUP in the peptide and in the identified protein are presented.

**Figure 7:**
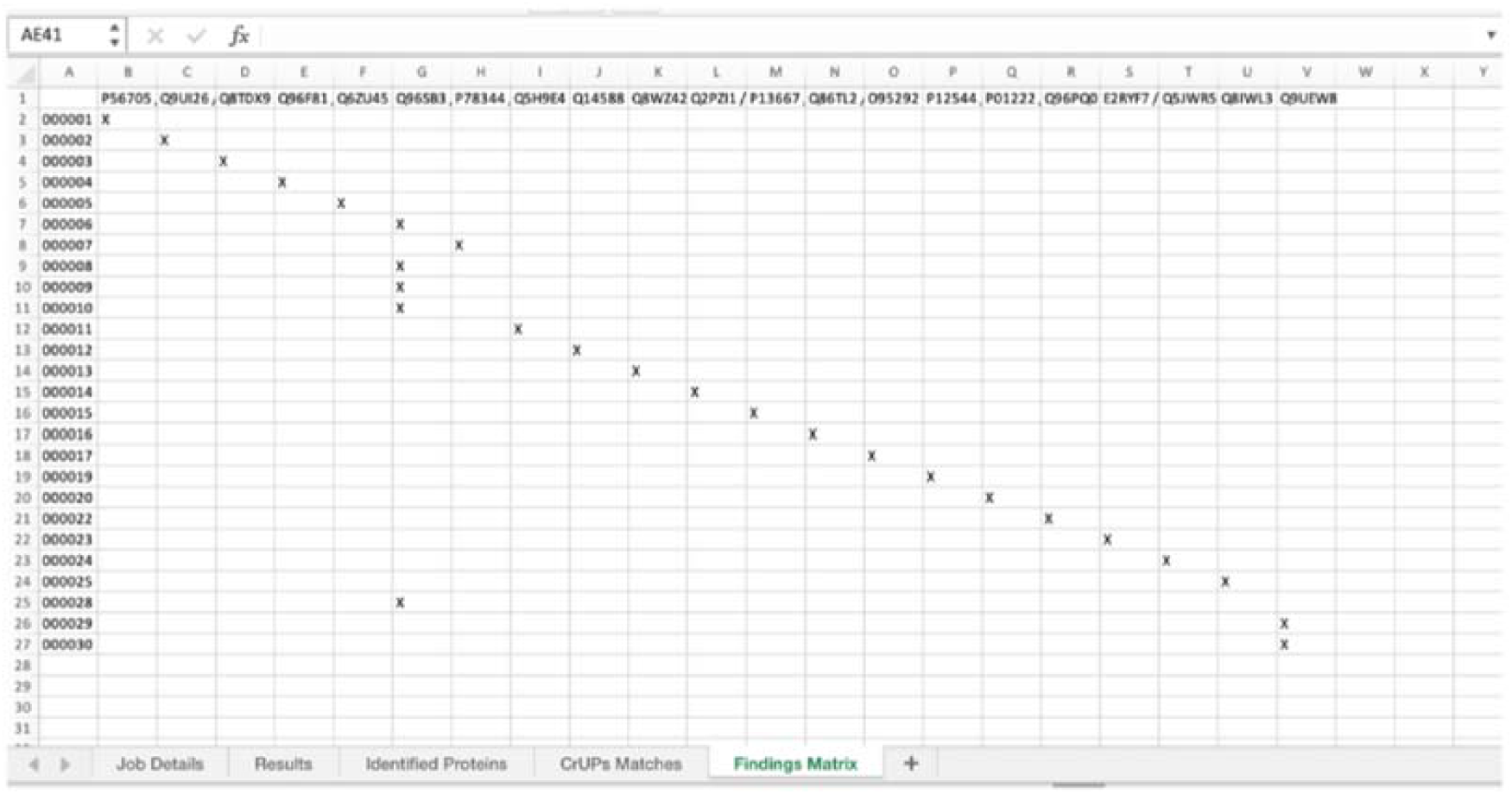
The “Findings Matrix” worksheet, displays all peptides that identify a protein, with an “X” marking the intersection between peptides and their corresponding proteins.

Finally, the *Findings Matrix* worksheet provides a global view of the results by displaying all peptides that identify a protein, with an “X” marking the intersection between each peptide and its corresponding protein.

### 3.4. UB-PIM notes

The UB-PIM approach contrasts with traditional protein identification methods, which search peptides within a FASTA format file containing theoretical peptides generated from protein digestion by a specified proteolytic enzyme. In conventional methods, if an identified peptide is present in the FASTA file, it is assigned to the protein from which it is theoretically derived (15).

According to the UB-PIM development concept, the method is independent of peptide type (e.g., tryptic digest peptides, chymotryptic digest peptides) and can be applied to any peptide. Based on this principle, UB-PIM is effective for protein identification in DIA procedures but is especially suitable for DDA procedures, where peptides are not predefined and only a subset of ions is selected for fragmentation during MS/MS. This indicates that UB-PIM enables secure protein identification in both bottom-up and top-down proteomic approaches.

Conventional proteomics guidelines require two unique peptides per protein to confirm an identification (16). While this rule minimizes false positives, it also excludes many biologically important but hard-to-detect proteins, such as small proteins, low-abundance proteins, and proteins with few proteotypic tryptic sites. These include biomarkers, tissue-specific proteins, hormones, and pathogen proteins, which are rare but crucial, and often difficult to detect and quantify.

By applying UB-PIM, even a single match to a CrUP can represent a high-confidence identification. This expands the detectable proteomic space and improves sensitivity without increasing false positive identifications. In this way, many biologically significant proteins (e.g., biomarkers, tissue-specific proteins, pathogen proteins) that yield only a few peptides under standard digestion—or are present at low abundance (e.g., signaling peptides, cytokines, membrane proteins)—can now be detected and quantified, overcoming the limitations of the two-peptide rule.

At the informatics level, the precomputed uniqueness of CrUPs simplifies post-processing of search results and enables instantaneous checking during peptide searches, resulting in faster and more secure identifications (17). Another major advantage of the approach is its straightforward integration into widely used search engines such as Proteome Discoverer, MaxQuant, MSFragger, and ProFound. Since CrUPs are highly specific and stable, they are also excellent candidates for targeted MS methods (SRM, PRM, DIA). In peptide identification, quantification, and targeted proteomics, shared peptides often cause signal contamination between proteins. The UB-PIM method avoids this entirely because of the uniqueness of CrUPs, ensuring that quantitation reflects true protein abundance rather than mixed contributions from homologues (17). This advantage is particularly evident in label-related methods, where UB-PIM could also support the design of ready-made peptide panels for validation and biomarker quantitation.

In this framework, it is not entire peptides but rather CrUPs within those peptides that are identified and quantified. This leads to rapid and secure identifications that accurately reflect protein abundance, shortens the pipeline from discovery to clinical or translational assays, and improves reproducibility. Instead of searching across an entire proteome database, smaller targeted databases containing only CrUP sequences (and their parent proteins) can be constructed—for example, databases restricted to cancer-related proteins. Such reduced databases minimize search space, memory usage, and false discovery rates, making them ideal for high-throughput clinical proteomics.

Finally, because CrUPs are derived from exact sequences, UB-PIM can distinguish isoforms, sequence variants, and post-translationally processed forms—an advantage particularly valuable for splice variant analysis or in differentiating highly similar paralogous proteins, which traditional peptide sets often cannot achieve. A future advancement of our method, currently in preparation, would involve transforming CrUPs into mass spectra (using the SEQUEST HT algorithm) and directly correlating CrUP-derived spectra with experimental mass spectra, thereby leveraging all the advantages outlined above (18).

## 4. Acknowledgments

This study has not been funded by any source whatsoever.

## 5. Authorship contribution

PV: Software, Methodology. KE: Validation, Software, Methodology, Formal analysis. SJD: Writing –review & editing, Writing – original draft, Validation. GThT: Conceptualization, Methodology, Writing –original draft, Writing – review & editing, Visualization, Validation, Supervision.

## Data and materials availability

All data are available in the main text and in the supplementary materials.

