## SUPPLEMENTARY MATERIAL for "A UNIQUOME BASED METHOD FOR THE PROTEIN IDENTIFICATION BY MASS SPECTROMETRY"

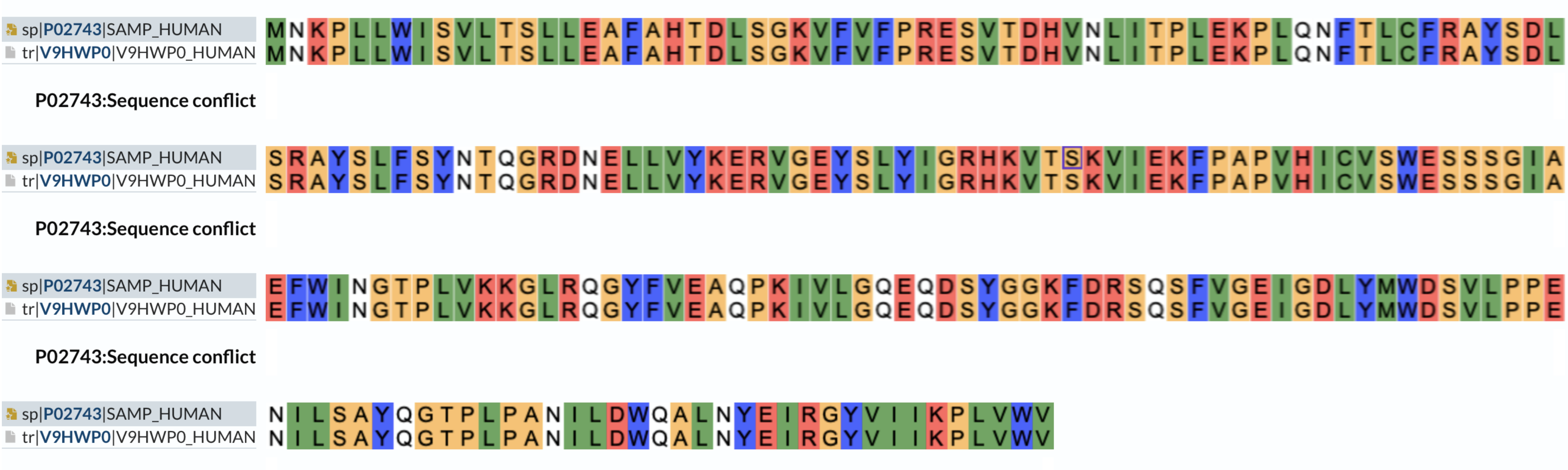


**Supplementary Figure 1.** Alignment of the SAMP_Human (P02743, Serum Amyloid P-component) and the V9HWP0_Human (V9HWP0, Pentraxin Family member) amino acid sequence. The sequence of these proteins are totally identical.

```
VW5B1_HUMAN     MPGLLNWITGAALPLTASDVTSCVSGYALGLTASLTYGNLEAQPFQGLFVYPLDECTTVI      60
E9PP07_HUMAN    MPGLLNWITGAALPLTASDVTSCVSGYALGLTASLTYGNLEAQPFQGLFVYPLDECTTVI      60
                ************************************************************

VW5B1_HUMAN     GFEAVIADRVVTVQIKDKAKLESGHFDASHVRSPTVTGNILQDGVSIAPHSCTPGKVTLD      120
E9PP07_HUMAN    GFEAVIADRVVTVQIKDKAKLE                                            82
                **********************

VW5B1_HUMAN     EDLERILFVANLGTIAPMENVTIFISTSSELPTLPSGAVRVLLPAVCAPTVPQFCTKSTG      180
E9PP07_HUMAN

VW5B1_HUMAN     TSNQQAQGKDRHCFGAWAPGSWNKLCLATLLNTEVSNPMEYEFNFQLEIRGPCLLAGVES      240
E9PP07_HUMAN

VW5B1_HUMAN     PTHEIRADAAPSARSAKSIIITLANKHTFDRPVEILIHPSEPHMPHVLIEKGDMTLGEFD      300
E9PP07_HUMAN

VW5B1_HUMAN     QHLKGRTDFIKGMKKKSRAERKTEIIRKRLHKDIPHHSVIMLNFCPDLQSVQPCLRKAHG      360
E9PP07_HUMAN

VW5B1_HUMAN     EFIFLIDRSSSMSGISMHRVKDAMLVALKSLMPACLFNIIGFGSTFKSLFPSSQTYSEDS      420
E9PP07_HUMAN

VW5B1_HUMAN     LAMACDDIQRMKADMGGTNILSPLKWVIRQPVHRGHPRLLFVITDGAVNNTGKVLELVRN      480
E9PP07_HUMAN

VW5B1_HUMAN     HAFSTRCYSFGIGPNVCHRLVKGLASVSEGSAELLMEGERLQPKMVKSLKKAMAPVLSDV      540
E9PP07_HUMAN

VW5B1_HUMAN     TVEWIFPETTEVLVSPVSASSLFPGERLVGYGIVCDASLHISNPRSDKRRRYSMLHSQES      600
E9PP07_HUMAN

VW5B1_HUMAN     GSSVFYHSQDDGPGLEGGDCAKNSGAPFILGQAKNARLASGDSTTKHDLNLSQRRRAYST      660
E9PP07_HUMAN

VW5B1_HUMAN     NQITNHKPLPRATMASDPMPAAKRYPLRKARLQDLTNQTSLDVQRWQIDLQPLLNSGQDL      720
E9PP07_HUMAN

VW5B1_HUMAN     NQGPKLRGPGARRPSLLPQGCQPFLPWGQETQAWSPVRERTSDSRSPGDLEPSHHPSAFE      780
E9PP07_HUMAN

VW5B1_HUMAN     TETSSDWDPPAESQERASPSRPATPAPVLGKALVKGLHDSQRLQWEVSFELGTPGPERGG      840
E9PP07_HUMAN

VW5B1_HUMAN     AQDADLWSETFHHLAARAIIRDFEQLAEREGEIEQGSNRRYQVSALHTSKACNIISKYTA      900
E9PP07_HUMAN

VW5B1_HUMAN     FVPVDVSKSRYLPTVVEYPNSAALRMLGSRALAQQWRGTSSGFGRPQTMLGEDSAPGNGK      960
E9PP07_HUMAN

VW5B1_HUMAN     FQALNMEASPTALFSEARSPGREKHGASEGPQRSLATNTLSSMKASENLFGSWLNLNKSR      1020
E9PP07_HUMAN

VW5B1_HUMAN     LLTRAAKGFLSKPLIKAVESTSGNQSFDYIPLVSLQLASGAFLLNEAFCEATHIPMEKLK      1080
E9PP07_HUMAN

VW5B1_HUMAN     WTSPFTCHRVSLTTRPSESKTPSPQLCTSSPPRHPSCDSFSLEPLAKGKLGLEPRAVVEH      1140
E9PP07_HUMAN

VW5B1_HUMAN     TGKLWATVVGLAWLEHSSASYFTEWELVAAKANSWLEQQEVPEGRTQGTLKAAARQLFVL      1200
E9PP07_HUMAN

VW5B1_HUMAN     LRHWDENLEFNMLCYNPNYV                                              1220
E9PP07_HUMAN
```

**Supplementary Figure 2.** The alignment of the reviewed protein VW5B1_Human (Q5TIE3, von Willebrand factor A domain-containing protein 5B1) and the unreviewed protein E9PP07_Human (E9PP07, von Willebrand factor A domain-containing 5B1), indicated that the E9PP07_Human is a fragment of VW5B1_Human protein.
